# Contingency Inverts Mammalian Herbivore Evolution in Australia

**DOI:** 10.64898/2026.03.04.709684

**Authors:** Aidan M. C. Couzens, Benedict King, Gavin J. Prideaux

## Abstract

The rise of Neogene herbivores with high-crowned (hypsodont) molar teeth has been viewed as a mostly predictable response to abrasive grazing diets. Using kangaroos, an isolated marsupial radiation, we show that the ancestral vertical slicing function of grazing kangaroo molars prompted heavy investment from the late Miocene in thickened enamel, rather than hypsodonty. Grazing kangaroo enamel thickness overlaps some robust hominins, evincing an eclectic, thick-enamelled grazer guild. The success of vertically-chewing marsupials contrasts with their placental counterparts, which were overwhelmingly replaced by transversely-chewing ungulates. This inversion is explained by the pre-grassland extinction of most transversely-chewing marsupials, and the crucial advent of thick enamel. These results challenge the determinism of the browser–grazer transition, and implicate extinction, and ensuing innovation, as causes of unpredictability in evolution.

## Main Text

The extent to which evolutionary outcomes are predictable due to the determinism of natural selection, or unpredictable due to the influence of chance events (historical contingency) has been much debated (*1–5*). Convergence between unrelated organisms that have evolved similar adaptations in order to exploit similar ecological niches has often been used to exemplify the deterministic nature of evolution (*5, 6*). However, much less attention has been directed toward explaining why organisms occupying similar ecological niches may fail to converge (*3*), or what role contingent factors like ancestry, extinction, innovation, or vicariance play in divergence versus convergence (*4*). Australia provides a unique theatre in which to examine these questions due to its long isolation (*7*) and unique radiation of marsupial mammals, which diversified into a range of terrestrial niches similar to those occupied by placental mammals elsewhere (*8*).

### Disparity and Divergence in Herbivore Evolution

Kangaroos (Macropodidae) are the most diverse and abundant mammalian herbivores on the Australian continent. Despite the peculiarity of their locomotory mode (as the only large-bodied hopping mammals), kangaroos have often been viewed as ecological analogues of artiodactyl ungulates, occupying a spectrum of dietary niches, with the larger, more derived macropodines filling grazing niches (*9–12*). Yet, the molar morphologies of grazing kangaroos and ungulates are highly disparate, reflecting distinct ancestries, and so the manner in which they chew abrasive grasses are different (*13*). Grazing ungulates (e.g., bovids, deer, horses) have selenodont molars with crescent-shaped crests that fracture food using a transverse shearing and grinding motion (*14*) (Fig. 1A). Although efficient mechanically, the cost is accelerated tooth wear due to the thin enamel needed to create a complex occlusal surface (*15*) (Fig. 1B). Ungulates countered this by evolving high-crowned, prismatic molars (*16–18*) (Fig. 1A), which maintain near-identical functionality throughout life (*14*). In contrast, grazing kangaroos have simpler, lower-crowned, bilophodont molars (*9*), dominated by two transverse blades or lophs (Fig. 1, A and B). These are adapted for cutting food in a mainly vertical motion, with only a minor transverse component (*13*). As the blades wear, kangaroo molars transition from cutting to crushing tools, with accelerated rates of blade destruction being a key indicator of their late Neogene dietary shift to grazing (*12*).

**Fig. 1.**
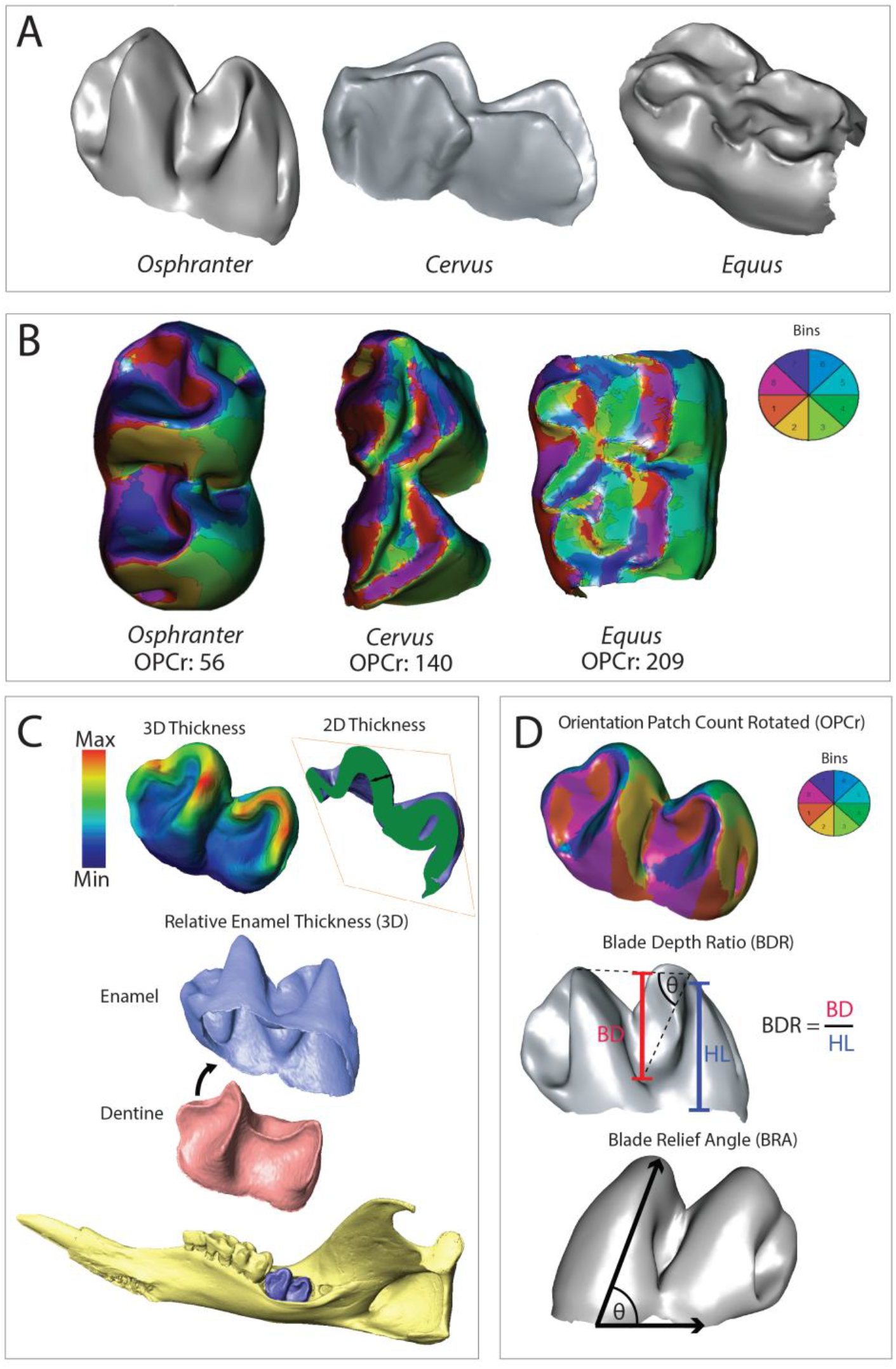
Macropod molar tooth shape and measurement of enamel thickness. (**A**) Comparison of orientation patch count (rotated) scores (OPCr) for a grazing macropodid (*Osphranter robustus*), the cervid *Cervus elaphus* and equid *Equus equus* x *caballus*. (**B**) Oblique views of lower molar structure. (**C**) Virtual extraction of enamel and dentine used to compute three-dimensional relative enamel thickness (RET3D), enamel thickness maps, and example of virtual 2D plane used to measure linear enamel thickness. (**D**) Measurements of OPCr), blade depth ratio, and blade relief angle. Scans for *Cervus* (ark:/87602/m4/M122164) and *Equus* (ark:/87602/m4/M105475) provided courtesy Idaho Museum of Natural History and downloaded from MorphoSource (www.MorphoSource.org).

Kangaroo molars resemble those of an eclectic range of evolutionarily waning or extinct vertically-chewing herbivore groups (*19*), including various proboscideans, some South American ‘native ungulates’, diprotodontoid marsupials, tapirs, manatees, and some apes and monkeys (e.g.,*17, 20–23*). But, by contrast, kangaroos experienced a profoundly different evolutionary fate, steadily increasing their diversity over the last 25 million years (Ma) (*10*). On most continents, the spread of grasslands and the rise of selenodont herbivores, especially artiodactyl ungulates, correlates with the decline of vertically-chewing placentals (*23–26*). These observations lead us to ask: A) How did kangaroos cope with abrasive grazing diets? B) Why did kangaroos never evolve ungulate-like molars? C) Among vertically-chewing herbivores, what drove the sustained success of marsupials, but the decline of placentals? This pattern is even more extraordinary given that the early Miocene rise of bilophodont marsupials coincided with the decline of several previously dominant selenodont marsupial groups.

### Thick Enamel as an Adaptation for Grazing

We used absorption X-ray microCT scanning to measure molar enamel thickness, a trait hypothesised to modulate resistance to hard or abrasive foods (*27, 28*). First, we confirmed that three-dimensional relative enamel thickness (RET3D), a measure of enamel thickness relative to dentine volume (see Methods), is an effective gauge of enamel investment relative to body mass. RET3D is a useful index of enamel thickness in fossil taxa where body mass is unknown. We find that dentine volume is highly correlated with body mass at the species level across Macropodoidea (R^2^ = 0.96) (fig. S1G). Molar enamel thickness, measured as RET3D or as volumetric enamel investment relative to body mass (mm^3^ _EVOL_/g _Bmass_ × 100), was 79% (18.3 ± 4.3 v. 10.3 ± 0.3) and 94% greater (0.60 ± 0.32 v. 0.31± 0.02), respectively, then in phalangerid and pseudocheirid possums, the closest extant marsupials to kangaroos (fig. S2A). Compared with primates, the only other mammalian group where enamel thickness has been comprehensively measured with microCT, median macropodoid enamel investment was 11% greater (ordinary least squares; OLS), or 22% greater (phylogenetic generalised least squares; PGLS) (fig. S3).

Overall, enamel investment relative to body mass in grazing kangaroos was 2.8 times greater than amongst browsers (1.09 ± 0.38 vs. 0.39 ± 0.24; PGLS _Enam. Invest. ∼ Diet_, DF=3, 9, F=3.632, p=0.06, Adj. R^2^=0.0.40; fig. S4C), and RET3D scores were 1.25 times greater amongst grazers compared with browsers (21.7 ± 3.2 vs. 17.4 ± 2.0; PGLS _RET3D ∼ Diet_, DF=3,9, F=4.4, p<0.05; Adj. R^2^=0.46; fig. S4B). Within these models, there were significant differences between grazers and browsers for both enamel investment (T value=3.21, p<0.05) and RET3D (T value=2.34, p<0.05). There were also significant differences between mixed feeders and browsers for enamel investment (T value=2.37, p<0.5) and RET3D (T value=3.62, p<0.01), and mixed feeders actually have higher RET3D (23.37 ± 4.0 vs. 17.44 ± 2.0; but not enamel investment) compared with grazers, suggesting that grass consumption rather than consumption level is important in driving enamel thickness variation. Most of the differences in enamel investment between grazers (or mixed feeders) and browsers appears to stem from thickened enamel along the lophid edge (PGLS _HCA ∼ Diet_, DF=3,9, p<0.05, Adj. R^2^=0.54). In grazers, enamel here is 1.5 times greater than in browsers (0.32 ± 0.09 vs. 0.22 ± 0.06; fig. S4A). Three-dimensional relative thickness maps show striking localisation of thick enamel along the crest edge in grazing and mixed-feeding kangaroos (fig. S2b). Although fungivorous kangaroos (potoroines, hypsiprymnodontids) have RET3D scores comparable to grazing kangaroos (21.5 ± 2.8 vs. 21.7 ± 3.2; fig. S5) their enamel investment relative to body mass and enamel thickness at the lophid edge are much lower (2.75, and 1.55 times, respectively). Additionally, in contrast to folivorous kangaroos, in fungivorous potoroines and hypsiprymnodontids the thickest enamel encircles the crushing basins (fig. S5). Rather than being indicative of abrasion resistance, the elevated RET3D of hypsiprymnodontids is mostly explained by their dentine reduced molars (fig. S1G), probably caused by reduction of their posterior molars (*29*). In potoroines, elevated RET3D could be indicative of hard-object feeding or abrasion caused by entrained grit associated with a more generalist diet (*11*).

Our results strongly suggest that thickened molar enamel in kangaroos is an adaptation to increased abrasive wear stemming from grazing. The link between thick enamel and dietary abrasion in kangaroos is consistent with evidence that phytolith abrasion rather than hard-object feeding (*30, 31*) explains thickened enamel in primates (*28*). Strikingly, the primates with the greatest enamel investment relative to body mass, the extinct robust hominins *Paranthropus robustus* and *Australopithecus africanus*, fall closest to grazing kangaroos like *Osphranter rufus* and *Macropus fuliginosus* (fig. S3). This bolsters the argument, hitherto based mainly on dental microwear and stable isotopes (*32*), that thick enamel in australopith hominins was an adaptation to resist abrasive wear from grasses or sedges (*33*).

Fossil data reveal parallel increases in enamel thickness in different lineages of bilophodont macropodids over the last 8 Ma (Fig. 2A). In lagostrophines and macropodines, measurements of linear enamel thickness show that greater enamel investment (Fig. 2B) is mostly correlated with thickening of enamel along the lophid (Fig. 2C). This was probably a mechanism to preserve cutting functionality as dietary abrasion escalated through the late Cenozoic as grass abundance increased in Australian ecosystems (*34, 35*). Amongst the proxies for enamel thickness (fig. S6, A to C), lophid-edge enamel thickness is the most strongly correlated with macrowear, a proxy for dietary abrasion (PGLS _HCA ∼Diet_, DF=1,16, F = 18.04, p<<0.01, Adj. R^2^ =0.50, fig. S6A). This is comparable to the correlation between crown height and macrowear (PGLS: DF=1,16, F=20.28, p<<0.01, Adj. R^2^=0.53, fig. S6D).

**Fig. 2.**
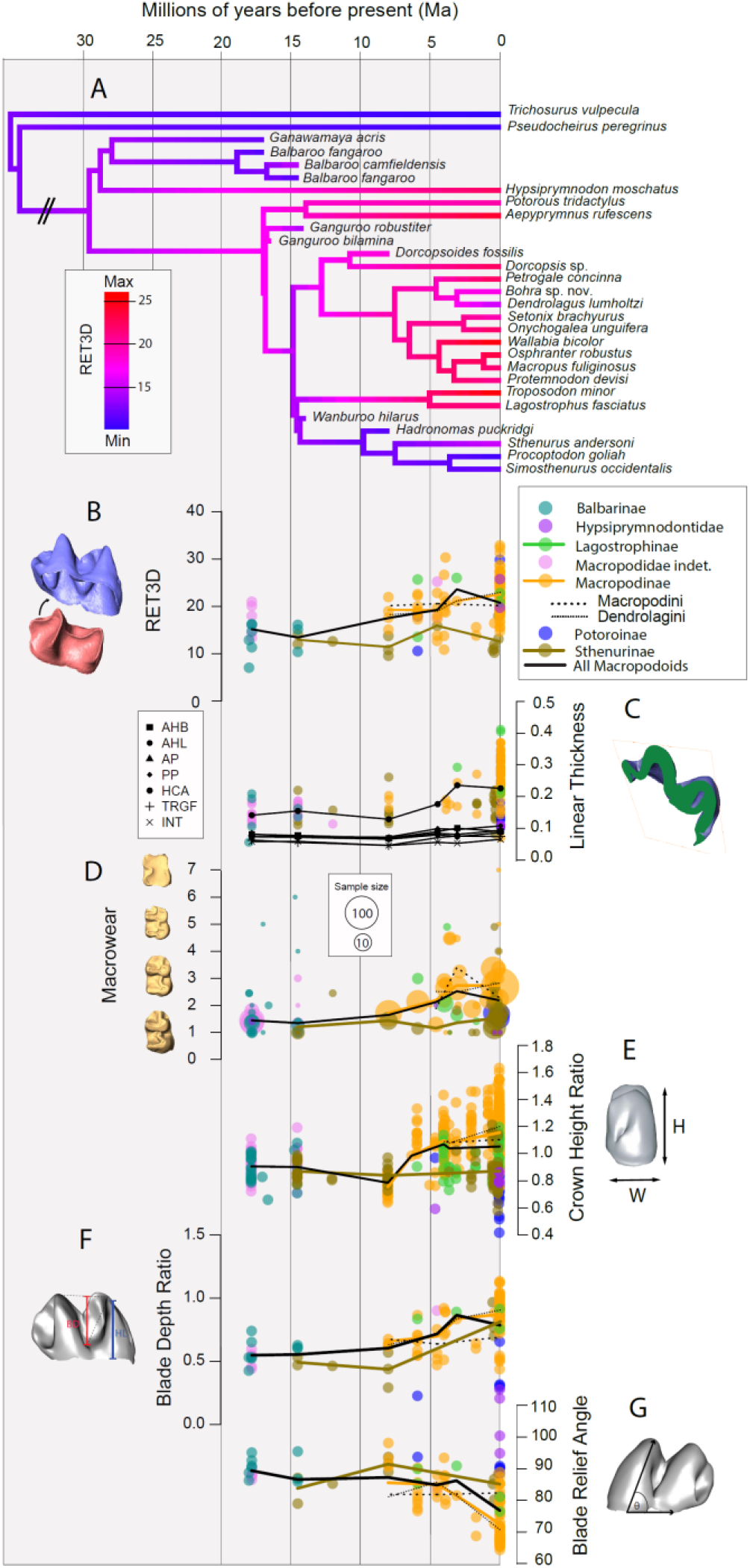
Macropod enamel thickness, dental wear, and tooth shape through time. (**A**) Phylogenetic reconstruction of three-dimensional relative enamel thickness (RET3D) across Macropodoidea. (**B**) Fossil RET3D over the past 20 million years (Ma). (**C**) Size-corrected linear enamel thickness across seven tooth regions. Abbreviations: anterior protolophid (AP), posterior protolophid (PP), trigonid fossa (TRGF), interlophid basin (INT), buccal face of anterior hypolophid (AHB), lingual face of anterior hypolophid (AHL), hypoconid apex (HCA). (**D**) Molar macrowear and (**E**) crown height ratios. (**F**) Molar blade depth ratio. (**G**) Blade relief angle.

Abrasion degrades the fracture efficiency of bladed teeth by causing opposing blades to misalign when fragmenting tough foods (*36*), akin to how worn scissors buckle rather than cut paper (fig.S7A). The recruitment of thick enamel was probably a mechanism to resist damage to the blade edge. Consistent with this inference, we find correlated shifts in enamel thickness (Fig. 2, B and C), lophid geometry (Fig. 2G) and dental macrowear (Fig. 2D). A vertical, ‘guillotine-like’ blade geometry (i.e., ∼90° blade relief angle, fig. S7A) characterises bilophodont macropodoids with low levels of macrowear (Fig. 2, D and G) like the thinly-enamelled Miocene balbarids, basal macropodids, and late Cenozoic sthenurines (Fig. 2, A to C). However, as enamel thickness and wear levels increased over the last 8 Ma (Fig. 2D), the lophids of lagostrophine and macropodine kangaroos became more anteriorly tilted (i.e., fig. S7, B and C), a shift we interpret as a further adaptation to maintain contact between the blade edges with increasing wear (Fig. 2G).

The initial increases in macropodid enamel thickness predate the mid-Pliocene rise of C_4_ grasses in Australia (*34*). Previous data suggested that elevated dental wear from c. 3.6 Ma drove increased crown height (*12*), but recent dating of the Curramulka Local Fauna to c. 6 Ma (*37*), an assemblage hitherto considered potentially Pliocene in age, shows that kangaroos were under pressure to evolve more durable molars before the end of the Miocene. This evidence predates the earliest palaeobotanical record of widespread C_4_ grasses (*34*), which dominate Australian grasslands today, by approximately 2.5 Ma. Recent pollen and biomarker data from offshore records suggest that C_4_ grass expansion in Australia may have been preceded by a C_3_ grass radiation 7–3.5 Ma ago (*35*). Together with elevated dust flux (*34*), which could have increased exogenous grit loads, we speculate that this early C_3_ phase was the key environmental driver of initial increases in macropodid enamel thickness.

### Contingent Evolution of Thick Molar Enamel

Rather than adopt an ungulate-like strategy of evolving very high crowned, occlusally complex molar teeth, our results show that kangaroos invested heavily in enamel thickness as an adaptation for grazing, with only very modest increases in crown height (*12*) (Fig. 2E). Even the highest-crowned grazing macropodid molars are more than 4-fold lower crowned than the highest-crowned ungulates (e.g.,*12, 16*). This led to the suggestion (*9*) that grazing macropodids were constrained to low-crowned molars by the limited range of transverse jaw movement permitted by their diprotodont mode of incisor occlusion. Yet, other diprotodont marsupials, including koalas and wombats, not only evolved significant transverse jaw movements, but ungulate-like selenodont molar morphologies (*13*).

Unlike the bunoselenodont ancestors of ruminant artiodactyls (*38*), the middle Miocene ancestors of grazing kangaroos had lost the stylar cusps and longitudinal ridge connecting the buccal upper molar cusps (*39*) present in the selenodont ancestor of diprotodont marsupials (*40, 41*). This likely diminished their potential to build an ungulate-like longitudinal loph or to broaden the upper molars to accommodate an extensive transverse occlusal stroke as occurs in ungulates (*9, 14*). Macropodids, show no evidence of evolving toward a low-relief (“prismatic”), ungulate-like molar structure; indeed, their blade depth shows a pronounced increasing trend over the last 8 Ma (Fig. 2F). This demonstrates increased adaptive investment in the bladed function of the molars. Instead, kangaroos retained bilophodont molars but acquired thickly-enamelled lophids in response to increased dietary abrasiveness, following an adaptive ‘path of least resistance’. Grazing macropodids were likely preadapted to this solution because basal macropodids already possessed moderately thick molar enamel (e.g., Fig. 2A), probably due to early Neogene recruitment of thickened enamel to form the lophid itself (*29*) (fig. S5). Balbarids, an early Neogene group of bilophodont macropodoids (*8*), lack thick enamel, especially along the crest edge (Fig. 2B, fig. S2, A and B), due to their peculiar dentine-based lophid structure (fig. S5). This may have placed them at an adaptive disadvantage as wear rates increased in the late Cenozoic (Fig. 2D), perhaps contributing to their extinction.

### Inverted Herbivore Evolution in Australia

The ascendency of grazing herbivores over the past 20 million years is one of the most striking trends in mammalian evolution (*26, 42*). In North America, Europe, and Asia, a period of early Paleogene experimentation was followed by a pronounced ‘winnowing’ of crown-type diversity that saw bunodont and lophodont herbivores replaced by those adapted for more transversely-directed modes of chewing like artiodactyls, rhinoceroses, and horses (*16, 42, 43*) (Fig. 3A). A similar succession occurred in isolated South America, where bilophodont xenungulates and pyrotheres were replaced by endemic, related lineages with higher crowned (*44*), more prismatic molars suited to transverse chewing (*23*). In Australia, this trend was inverted. From the late Oligocene, the species richness of selenodont vombatiforms (ilariids, wynyardiids, mukupirnids) shows a striking decline, being overtaken by increasingly diverse vertically-chewing macropodoids and diprotodontoids (*8*) (Fig. 3B). This transition was part of a broader faunal turnover that saw other ‘archaic’ marsupial groups (e.g., balbarids, miralinids, yalkaparidontids, yaralids) replaced by more modern clades (*8, 12, 45, 46*).

**Fig. 3.**
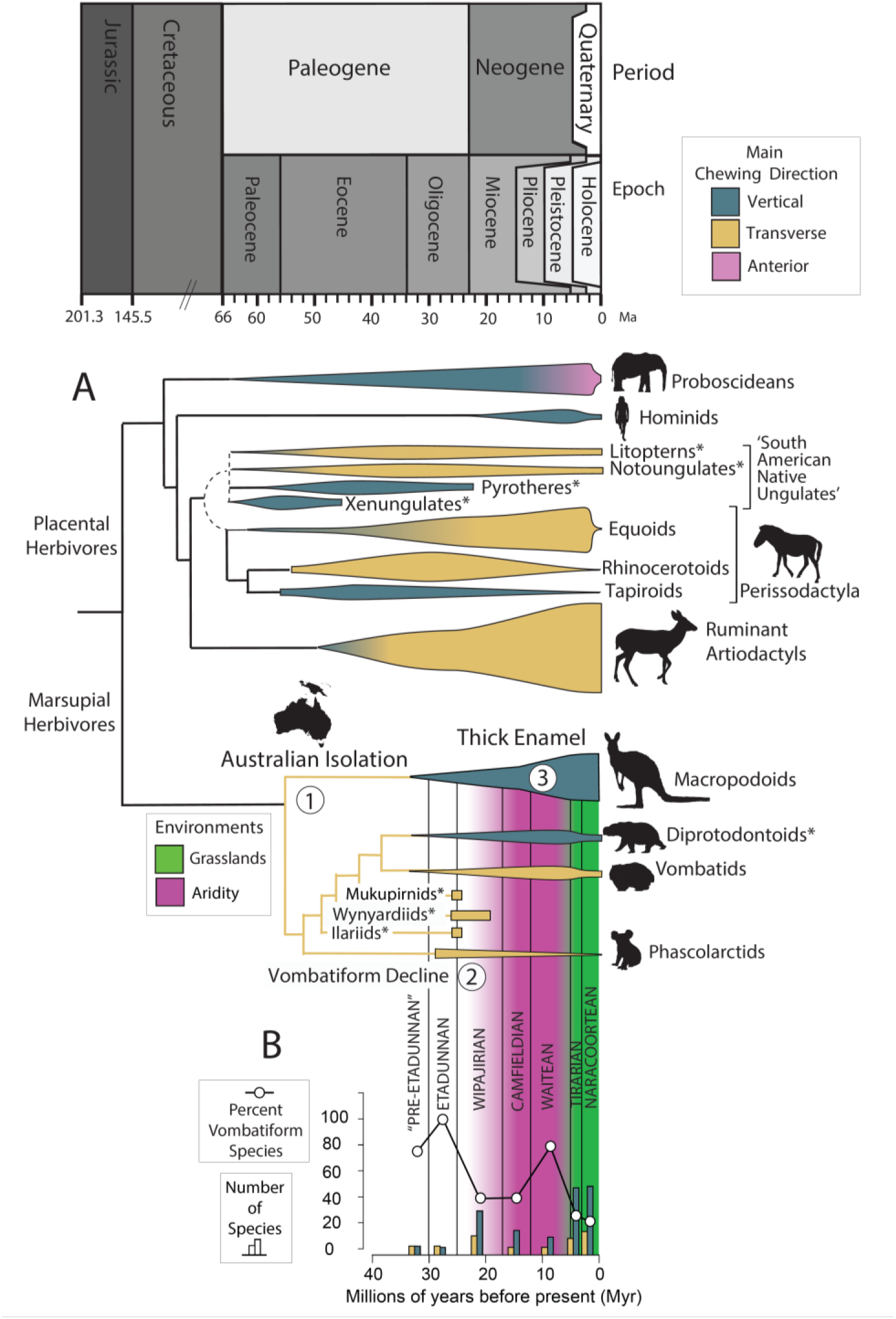
Divergent histories of marsupial and placental herbivores. (**A**) Patterns of herbivore clade richness relative to chewing direction. Following Australian isolation in the late Eocene (‘1’), marsupials diversified into vertically- and transversely-chewing clades dominated by the macropodids and diprotodontoids, and vombatiforms, respectively. (**B**) From the early Neogene, vombatiform richness declines (‘2’) due to extinction of basal transversely-chewing lineages (e.g., ilariids, wynardiids, mukupirnids). Vertically-chewing macropodoids and diprotodontoids diversify as Australian ecosystems became more arid, with some macropodoids acquiring thick molar enamel in the late Miocene (‘3’), facilitating expansion into a grazing niche. Australian land mammal ages follow (*58*). Vertical width of clades approximates generic richness.

Thick enamel was a key innovation for kangaroos, but it only partly explains the inverted herbivore history of Australia, because other successful vertically-chewing lineages, like the diprotodontoids, had molar enamel less than half that of kangaroos (RET3D: 7.8 ± 0.9 vs. 18.3 ± 4.3; fig. S2). In addition, several vertically-chewing placentals, such as *Paranthropus boisei* (*47*) and ‘gomphothere’ proboscideans (*48*), possessed unusually thick enamel, but nevertheless became extinct. In the case of *P. boisei*, dental microwear and stable-carbon isotopes show that it shared a monocot diet with pigs, hippopotamuses, horses and bovids shortly before its extinction (*49*). Similarly, late Miocene gomphotheres were increasing the proportion of grass in their diet, as well as their enamel thickness, when replaced by thinner-enamelled grazing ruminants and elephants (*24, 48*). The repeated advent of thick enamel amongst widely divergent herbivore groups suggests that this might be the most evolvable of a limited number of adaptive pathways open to vertically-chewing herbivores that shift to grazing.

The decline of selenodont herbivores in Australia may have been linked to a suite of pre-existing digestive and locomotory traits which disadvantaged them in drying environments. In the arid, nutrient-poor grassland, shrubland and desert now typical of much of Australia (*50*), hindgut-fermenting vombatiforms were probably less efficient at extracting and retaining nutrients than foregut-fermenting kangaroos (*51*). Additionally, the squat body plan of early terrestrial vombatiforms (*8, 52*) was potentially adaptively constraining, perhaps tethering them to drying water sources, whereas more vagile kangaroos could relocate. By enabling wombats to better retain water, fossoriality was probably a key reason for their comparative success (*53*). Several other burrowing or crevice-dwelling marsupial lineages also prospered in arid ecosystems, including several dasyurid lineages, marsupial moles, bilbies, bettongs (*45*) and, post-European arrival, introduced rabbits (*54*). Diprotodontoids probably benefited from the improved water retention, nutrient extraction, and thermal tolerance associated with their gigantism (*52*).

We attribute the unusual dominance of vertically-chewing grazers in Australia to three main events: 1) long geographic isolation, 2) the extinction of most selenodont forms before the spread of Australian grasslands; and 3) the appearance and fixation of mutations for thick enamel (Fig. 3). Save for a single tooth from the 55-Ma-old Tingamarra Local Fauna, interpreted as a condylarth (*55*), placental herbivores are not thought to have reached Australia until European arrival. Australia’s geographic isolation probably benefited vertically-chewing marsupials by preventing the arrival of cursorial, foregut-fermenting competitors. If ruminants had have reached Australia before the mid-Cenozoic arid shift, it is plausible that macropodids could have suffered declines, like other thick-enamelled herbivores. Notably, though, the failure of introduced ungulates to outcompete grazing macropodids (*56*) suggests that such an outcome would be far from assured, perhaps due to the closely-matched locomotory and digestive traits of these two groups (*10, 11, 57*). Indeed, the unique success of vertically-chewing marsupials in Australia suggests that locomotory and physiological traits may be far more pivotal than dental traits in determining the fates of herbivore lineages. The extinction of most selenodont vombatiforms 7–10 Ma before the spread of grasslands, left Australia curiously dominated by vertically-chewing herbivores. By ‘reshuffling the deck of cards’ in evolution, extinctions, and the ensuing innovation they enable, can scuttle forecastable outcomes, perhaps explaining why the predictability of evolution appears to decay at longer timescales (*1, 3, 4*).

## Acknowledgments

For scanning and technical assistance, we thank M. Skinner, T. Senden, M. Turner, J.-J. Hublin, P. Schöenfeld, H. Temming, W. Handley, A. Limaye, B. Viola and C. Moore. Collections access and information was provided by M.-A. Binnie, K. Butler, L. Dawson, S. Hocknull, S. Ingelby, H. Janetzki, R. Lawrence, J. Louys, R. Palmer, D. Pickering, P. Murray, A. Rozefelds, K. Spring, D. Stemmer, K. Travouillon, L. Umbrello, J. Wilkinson, A. Yates. T. Ziegler, and Y.Y. Zhen. R. Wells, M. Skinner, S. Arman, G. Gully, A. Crichton, and H. Zhang are thanked for discussions and/or feedback.

## Author contributions

Conceptualization: GJP, AMCC.

Data Collection: AMCC

Analysis: AMCC, BK

Writing: AMCC, GJP, BK

## Supplementary Materials

Materials and Methods

Figs. S1 to S8

References (59–79)

