## Supplementary information for "Contingency Inverts Mammalian Herbivore Evolution in Australia"

**The PDF file includes:**

Materials and Methods
Figs. S1 to S8
References #59–79.

### Materials and Methods

#### Scanning

Mandibular molars from fossil and modern macropods were sourced from the Australian Museum, Australian National Wildlife Collection, Flinders Palaeontology, Queensland Museum, South Australian Museum, Museums Victoria and Western Australian Museum. Specimens were scanned using high resolution X-ray absorption microcomputed tomography (microCT) at the following facilities: Department of Applied Physics, Australian National University (custom helical microCT), Flinders Microscopy (Nikon XTH225ST), Max Planck Institute for Evolutionary Anthropology (Actis BIR 225 industrial microCT, SkyScan 1072), and University of Adelaide Microscopy (SkyScan 1076). Specimens were scanned at isometric voxel-sizes ranging from 8.6–41µm.

#### Linear Enamel Thickness

Linear enamel thickness was measured from seven tooth regions with the *oblique slice* tool in Avizo 8 (fig. S8). Due to large variations in crown morphology between macropodoids, with the exception of cusp tip measurements, we avoided strictly defining section planes from landmarks. Instead, section planes were positioned across functionally homologous regions which displayed locally homogenous enamel thickness (cf. *59*). Oblique sections relative to the enamel-dentine junction (EDJ) can overestimate enamel thickness (*60*), so we report the shortest distance from the EDJ to outer-enamel surface (OES). The half maximum H method (*61*) was used to define material boundaries. The seven linear measurements were obtained (fig. S1) which included: anterior protolophid (AP), posterior protolophid (PP), floor of the trigonid fossa (TRGF), interlophid basin (INT), buccal face of the anterior hypolophid (AHB), lingual face of the anterior hypolophid (AHL), and hypoconid apex (HCA). Except for HCA, where the plane was defined explicitly, error in section plane orientation was accommodated by three repeat measurements. Linear thickness measurements were standardised for body size by computing a ratio relative to maximum width of the trigonid or talonid, which is highly log-correlated with body mass ($R^2$= 0.95; fig. S1,C and D).

#### Three-Dimensional Enamel Thickness

To measure dental tissue volume and compute RET3D, teeth were digitally segmented with Avizo 8. Raw image stacks were filtered with a mean of least variance filter implemented with MIA software (*62*). The filtered volumes were segmented into ‘enamel’, ‘dentine’, and ‘air’ labels using ‘Edit new label field’ function. Resulting segmentations were assessed manually (slice by slice) against unfiltered volumes and adjustments made if needed. A surface model of the enamel cap was created with the ‘Generate surface’ module. Enamel volume was measured using the ‘Surface area volume’ function. To measure EDJ surface area and dentine volume of the enamel cap, the EDJ was extracted and flat-filled using the ‘Fill holes’ function in Geomagic studio 12. Both the EDJ surface and the flat-filled EDJ cap were reimported into Avizo to measure the surface area and volume, respectively. RET3D was calculated following (*47*) as:

$$\mathrm{RET3D} = 100 \times \frac{(\mathrm{EVOL}/\mathrm{EDJSA})}{\sqrt[3]{\mathrm{DVOL}}}$$

RET3D = Three–dimensional relative enamel thickness

EVOL = Enamel volume (mm$^3$)

EDJSA = Surface Area of enamel–dentine junction (mm$^2$)

DVOL = Dentine volume of the enamel cap (mm$^3$)

To characterise variation in enamel thicknesses across the tooth surface we extracted the OES. Using the 'Surface distance' function a surface distance matrix was computed measuring shortest distance from OES to EDJ. The 'Surface view' module was used to visualise the distance matrix on the OES.

Tooth Shape Measurements

Three aspects of tooth shape were measured: blade relief angle, blade depth ratio, and orientation patch count rotated (Fig.1D). Blade relief angle describes the anterior inclination of the lophid, measured as the angle between the hypoconid apex relative to a line bisecting the buccal most extent of the enamel cervix on the hypolophid and protolophid. Blade depth ratio is an index of the depth of the interlophid valley relative to the lingual crown height at the hypolophid. This was computed by measuring distances corresponding to a right-angle triangle: the distance between the interlophid valley floor and the centre of the hypolophid crest (hypotenuse) and the half distance connecting the centre of the protolophid and hypolophid crests (adjacent). Next, the angle connecting the minimum of the interlophid valley, the centre of the hypolophid crest, and the centre of the protolophid crest was determined. Using the Law of Sines the blade depth (opposite side) was computed this divided by the lingual crown height of the hypolophid gives the blade depth ratio. Tooth complexity was measured with orientation patch count rotated (OPCr) (*63, 64*). The surface file of the outer enamel surface was oriented in Avizo 8, smoothed in Geomagic studio 12, and triangle count reduced to 10,000 faces with the 'vcgQEdecim' function in the R package Rvcg (65). The OPCr value of the resulting surface was calculated with 'molaR_batch' in the R package molaR (*66*).

Phylogenetic and Data Analysis

Data on primate enamel thickness was drawn from *(47,67,68)*. Body mass estimates for macropodids and primates were drawn from (*69*–71). For phylogenetic analysis of diet and enamel thickness we used the molecular phylogeney of extant marsupials from (*45*). For joint analysis of fossil and modern enamel thickness we used a modified version of the the node and tip dated total evidence tree for macropodoids from (*72*), and for primates, the consensus time-tree of (*73*). We modified the macropodoid phylogeny by grafting key fossil taxa on to this backbone tree using the 'tree.merger' function in the package 'Rrphylo' (*74*). Phylogenetic relationships and tip ages for balbarine kangaroos were based on (*75*). We then recalibrated node ages along the stem using the 'scaleTree' function using molecular divergence ages from (*45*) for five splits: *Hypsiprymnodon–Trichosurus*, *Hypsiprymnodon–Potorous*, *Lagostrophus–Potorous*, *Lagostrophus–Macropus*, and *Macropus–Dendrolagus*. Tip species lacking data were replaced with congenerics with data coverage. Ancestral state reconstruction was performed using the packages 'ape' (*76)* and 'phytools' (*77*). Phylogenetic generalised least squares regression was performed using the pgls function in the package 'caper' (*78*). To analyse marsupial herbivore diversity through time we extracted fossil occurences from the occurrence matrix of (*58*). We computed the proportion of total marsupial herbivore species which were vomabtiforms in each Australian Land Mammal Age (ALMA) defined by (*58*) and plotted the mid-point age for each

ALMA. Next we tallyed the number of species in each ALMA represented by herbivores with different chewing modalities. Herbivores with ‘bunodont’, ‘bilophodont’, or ‘bunolophodont’ molars were classifed as ‘vertically-chewing’ and those with ‘selenodont’, ‘bunoselenodont’ or ‘selenolophodont’ molars as ‘transversely-chewing’. Data analysis was performed using the R programming language (version 4.2.2) (*79*). For all results, means are reported plus or minus one standard deviation and an alpha level of 0.05 was used for statistical significance.

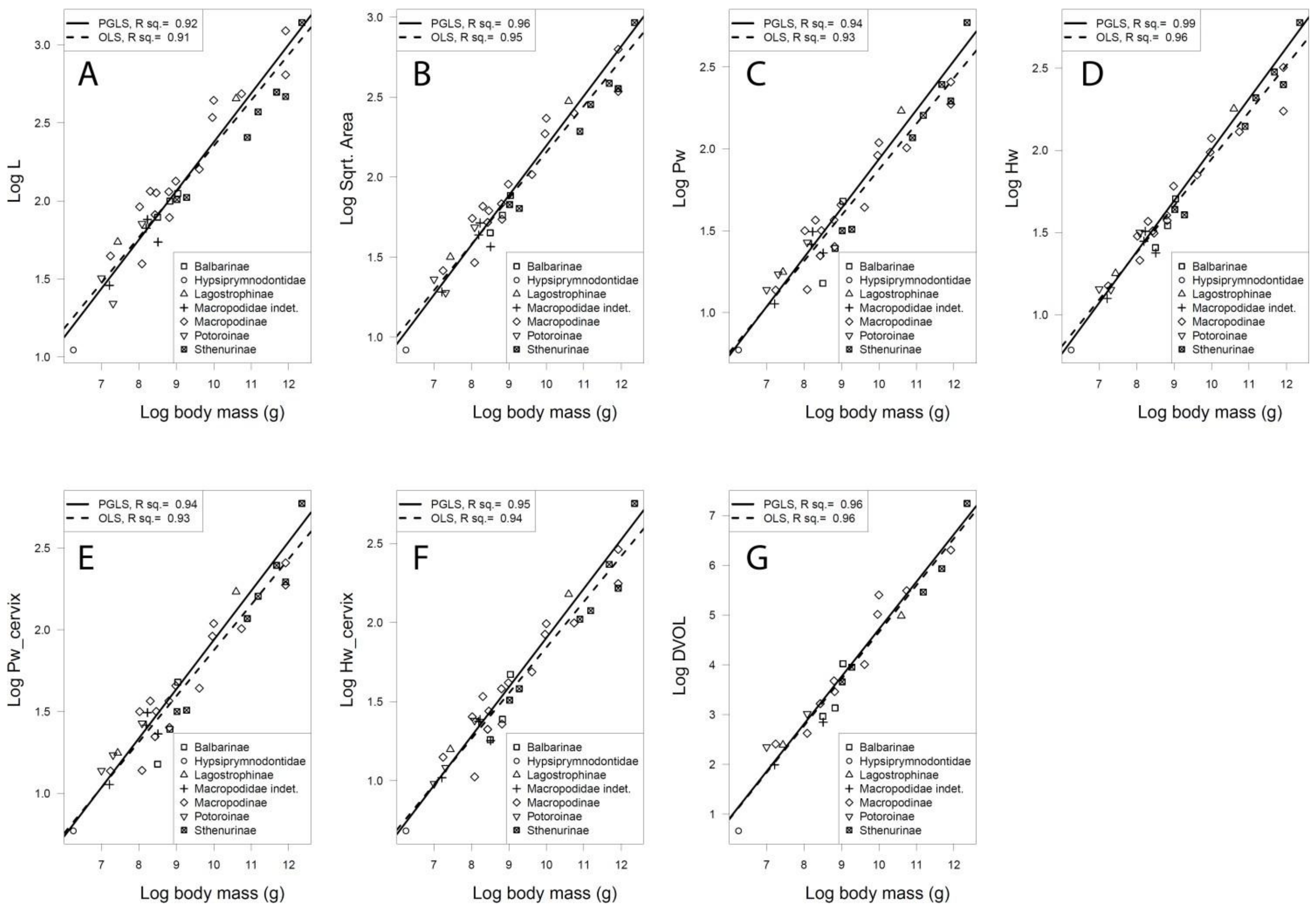


**Fig. S1. Correlations between dental measurements and body mass across macropodoid species.** (**A**) Log molar length. (**B**) Log square root of molar area. (**C**) Log maximum protolophid molar width. (**D**) Log maximum hypolophid molar width. (**E**) Log protolophid molar width at the enamel cervix. (**F**) Log hypolophid molar width at the enamel cervix. (**G**) Log dentine volume of the enamel cap. Solid lines is the phylogenetic generalised least squares (PGLS) regression best fit. Dashed line is ordinary least squares (OLS) regression best fit.

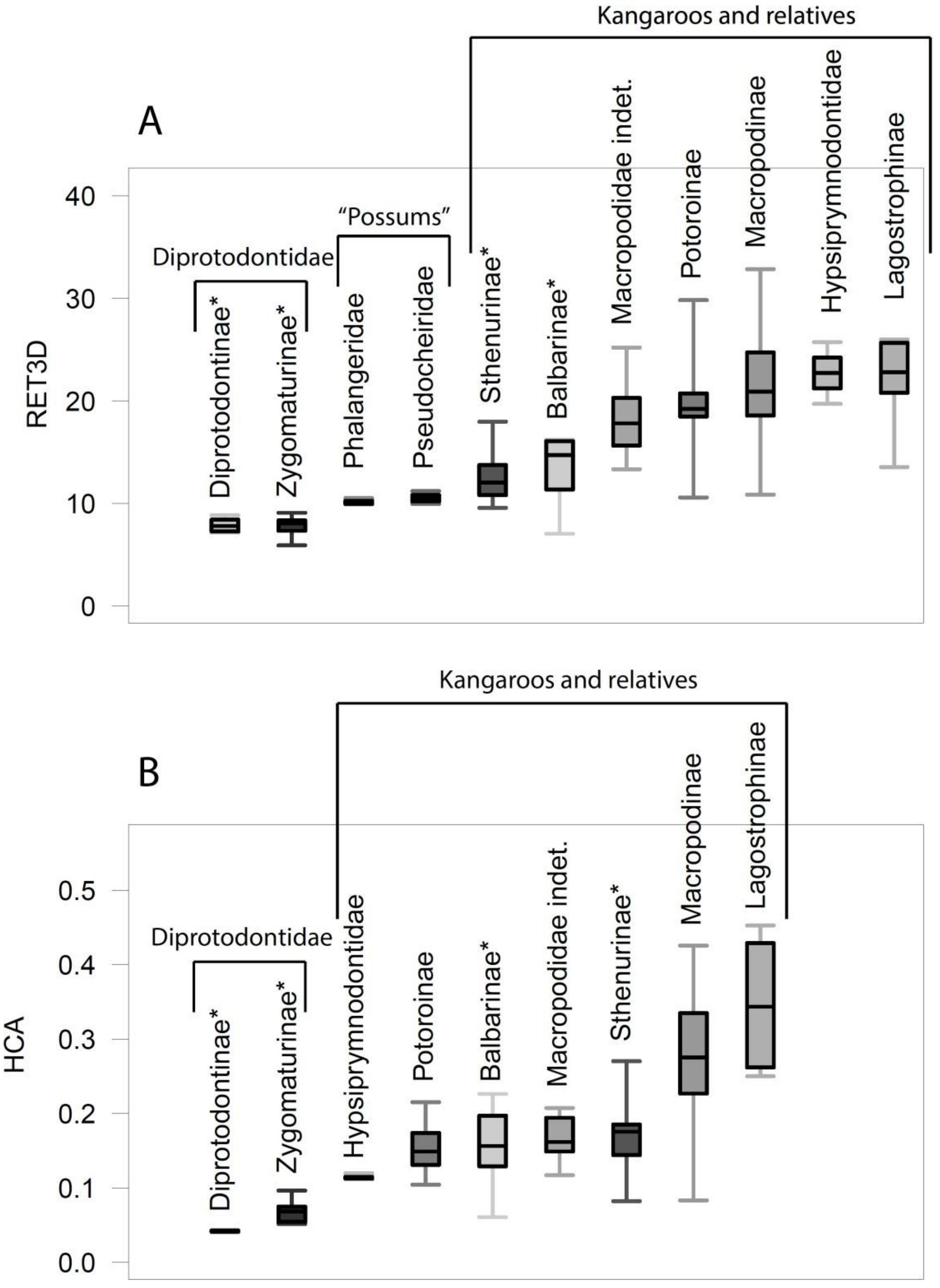


**Fig. S2. Box and whisker plot of clade-level variation in macropodoid enamel thickness.** (**A**) Three-dimensional relative enamel thickness (RET3D). (**B**) Linear enamel thickness at the hypoconid cusp apex (HCA). Asterix denotes wholly extinct clades.

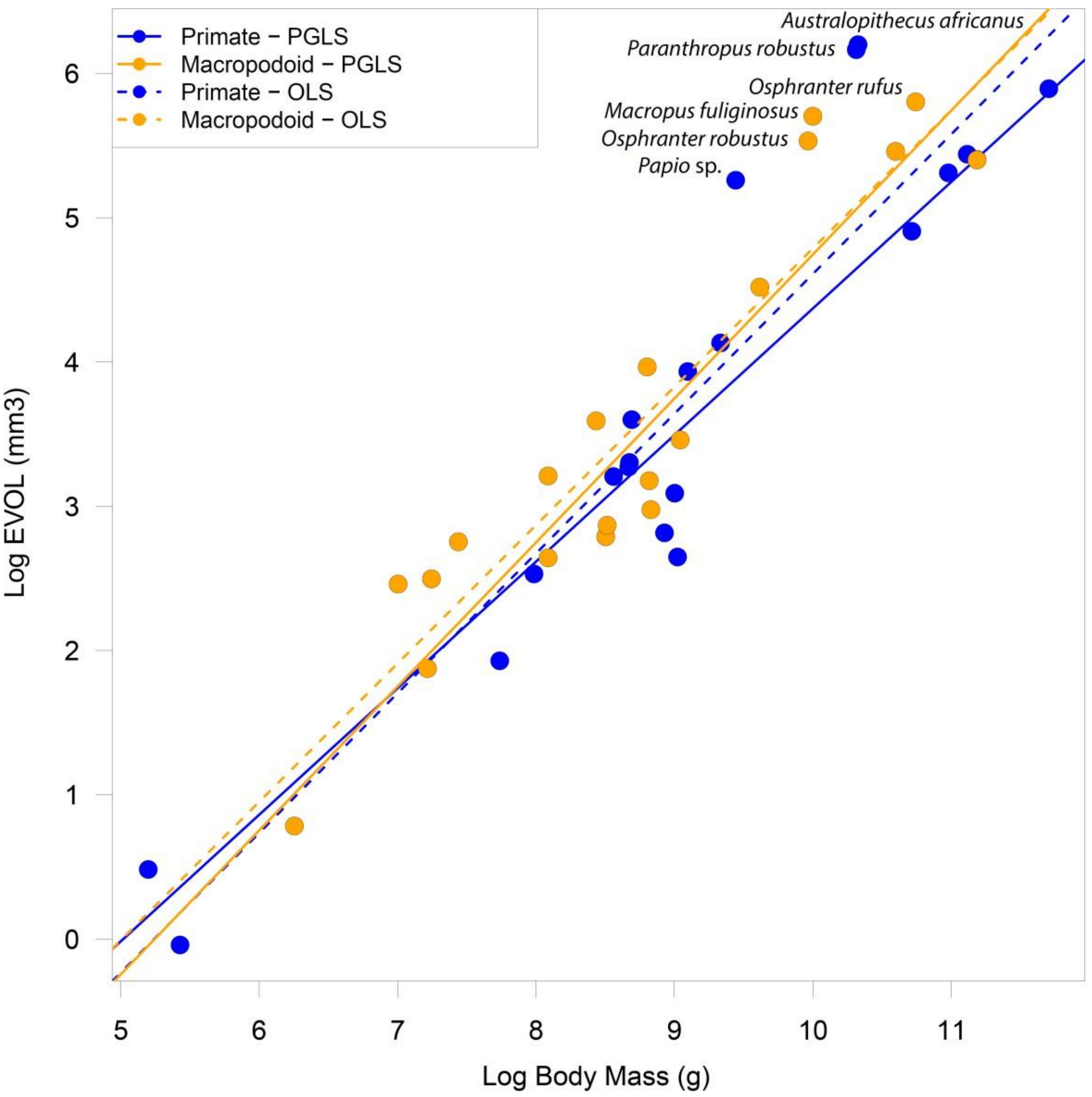


**Fig. S3. Enamel investment relative to body mass in macropodoids and primates.** Log–log major axis regression of molar enamel volume (mm$^3$) against body mass (g) in 21 species of fossil and modern primate (blue points) and 36 species of fossil and modern macropod (orange points). Solid lines are phylogenetic generalised least squares regression (PGLS). Dashed lines are ordinary least squares regression (OLS).

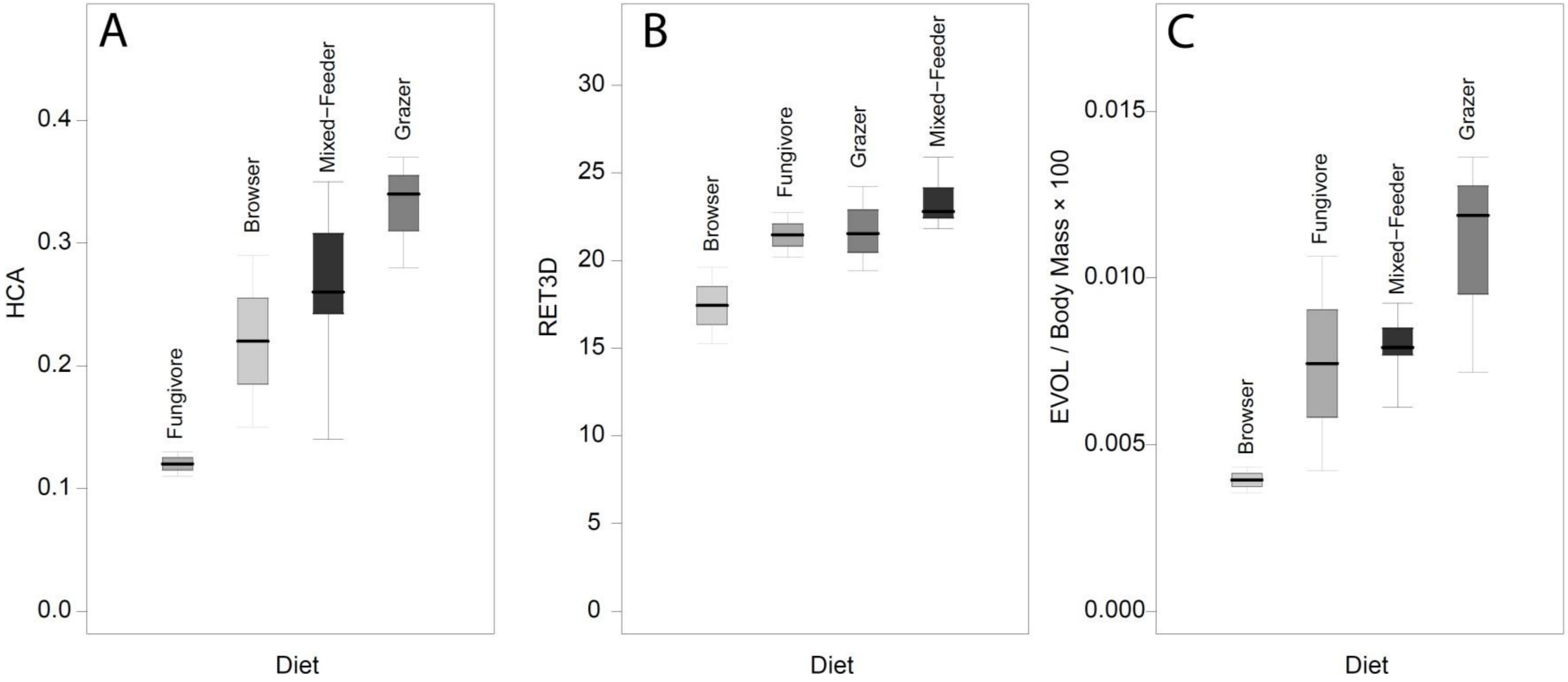


**Fig. S4. Box and whisker plot of variation in enamel thickness amongst extant macropodoid species for diet groupings.** (**A**) Linear enamel thickness at the hypoconid cusp apex (HCA). (**B**) Three-dimensional relative enamel thickness (RET3D). (**C**) Enamel volume relative to body mass (g) (× 100).

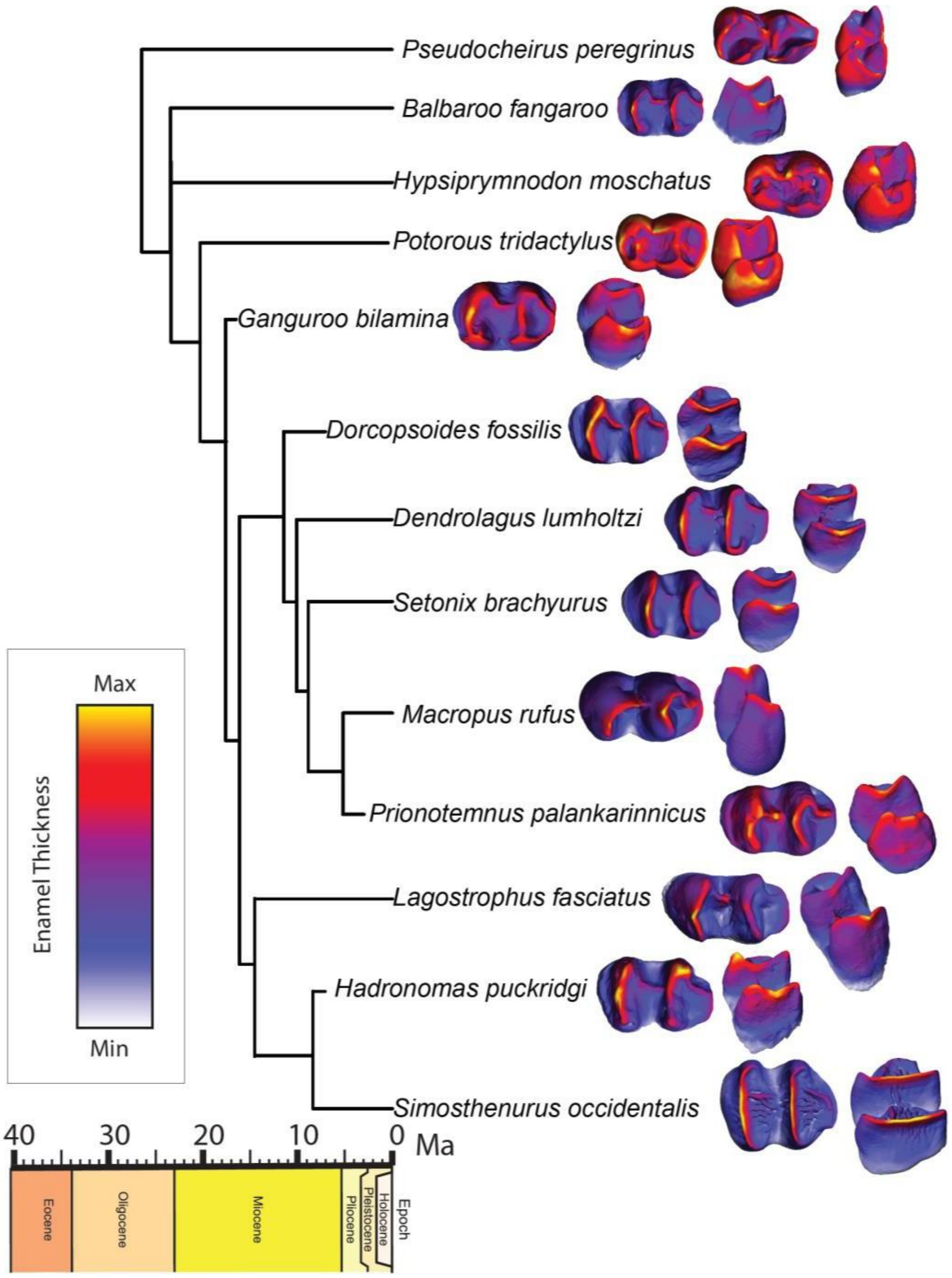


**Fig. S5. Relative lower molar enamel thickness in fossil and living macropodoids.** Occlusal and oblique posterobuccal views. Teeth not to scale.

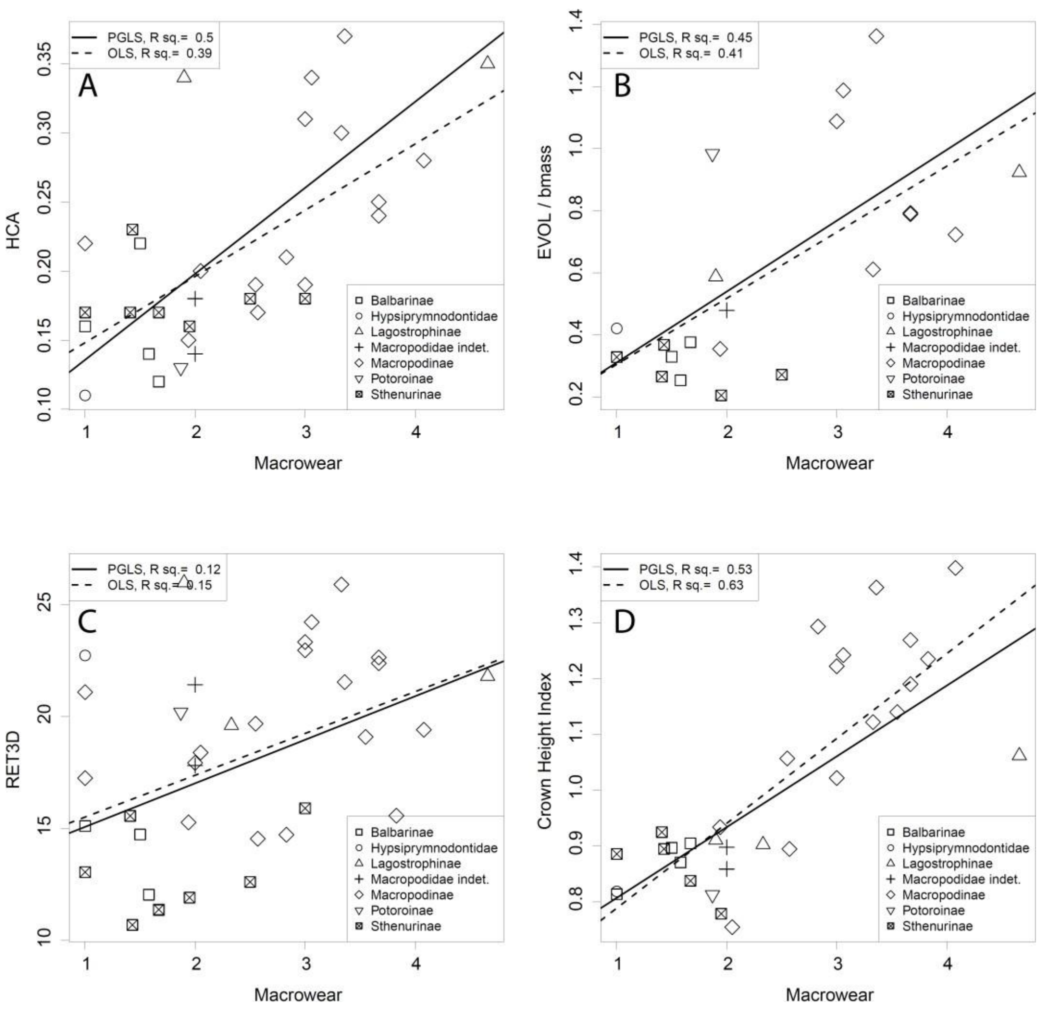


**Fig. S6. Regressions of dental traits against molar macrowear for fossil and modern macropodoid species.** (**A**) Enamel thickness of the hypoconid cusp apex (HCA). (**B**) Ratio of molar enamel volume ($mm^3$) to body mass (g) (×100). (**C**) Three-dimensional relative enamel thickness (RET3D). (**D**) Crown height index. Solid lines are phylogenetic generalised least squares regressions (PGLS). Dashed lines are ordinary least squares regression (OLS).

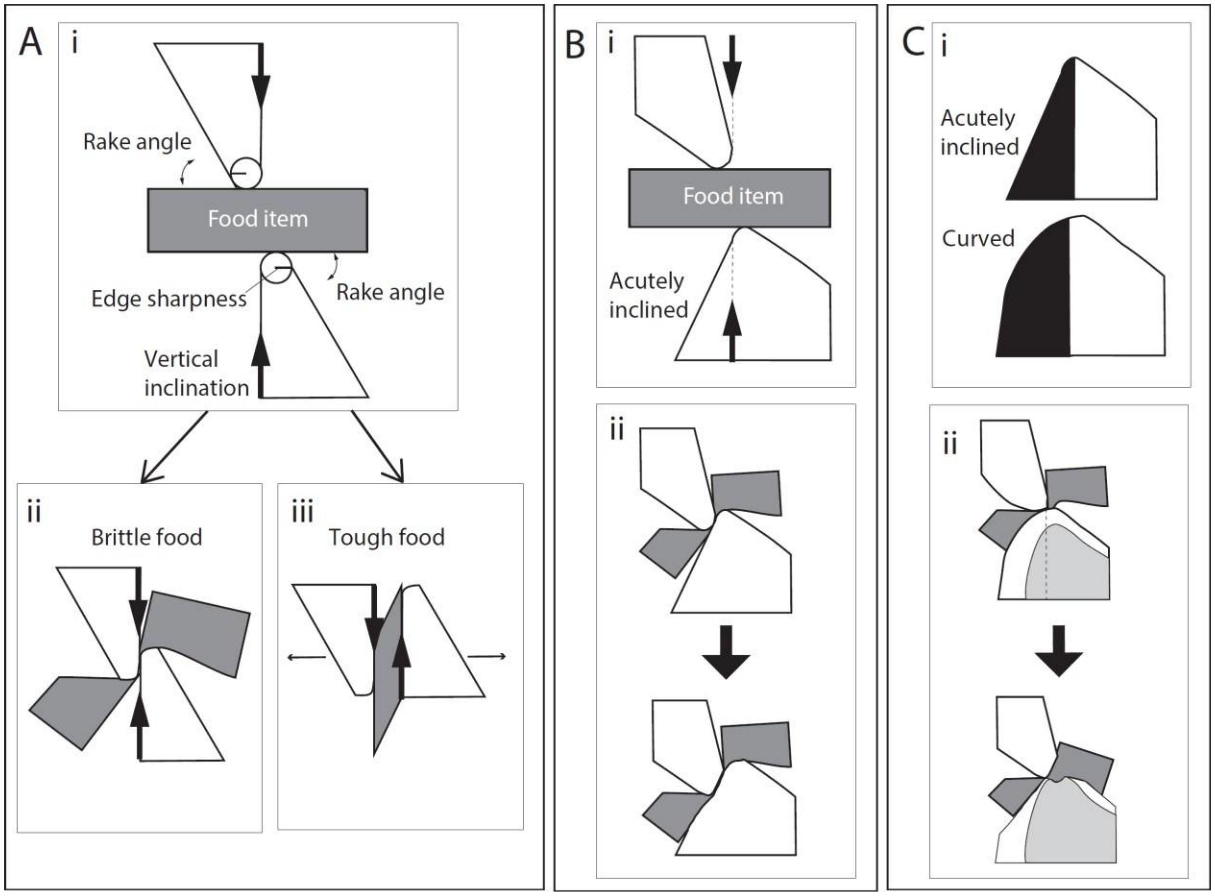


**Fig. S7. Dynamics of food fracture in a bladed system subject to dietary wear.** (**A**) (i) Schematic transverse view of food fracture between vertically-oriented blades modified from *(36)*. (ii) Complete fracture of brittle food. (iii) Incomplete fracture of food with high toughness due to blade separation. (**B**) (i) Food fracture with inclined relief surface leads keeps blades in contact and increases blade volume behind point of contact (ii). Formation of a wear land disperses fracture force leading to incomplete fracture. (**C**) (i) Curvature and inclination of the relief surface increases further blade volume. (ii) Durability of blade edge increased with thick enamel. With blade wear, preferential dentine erosion facilitates capture and fracture of food items at the exposed enamel–dentine junction.

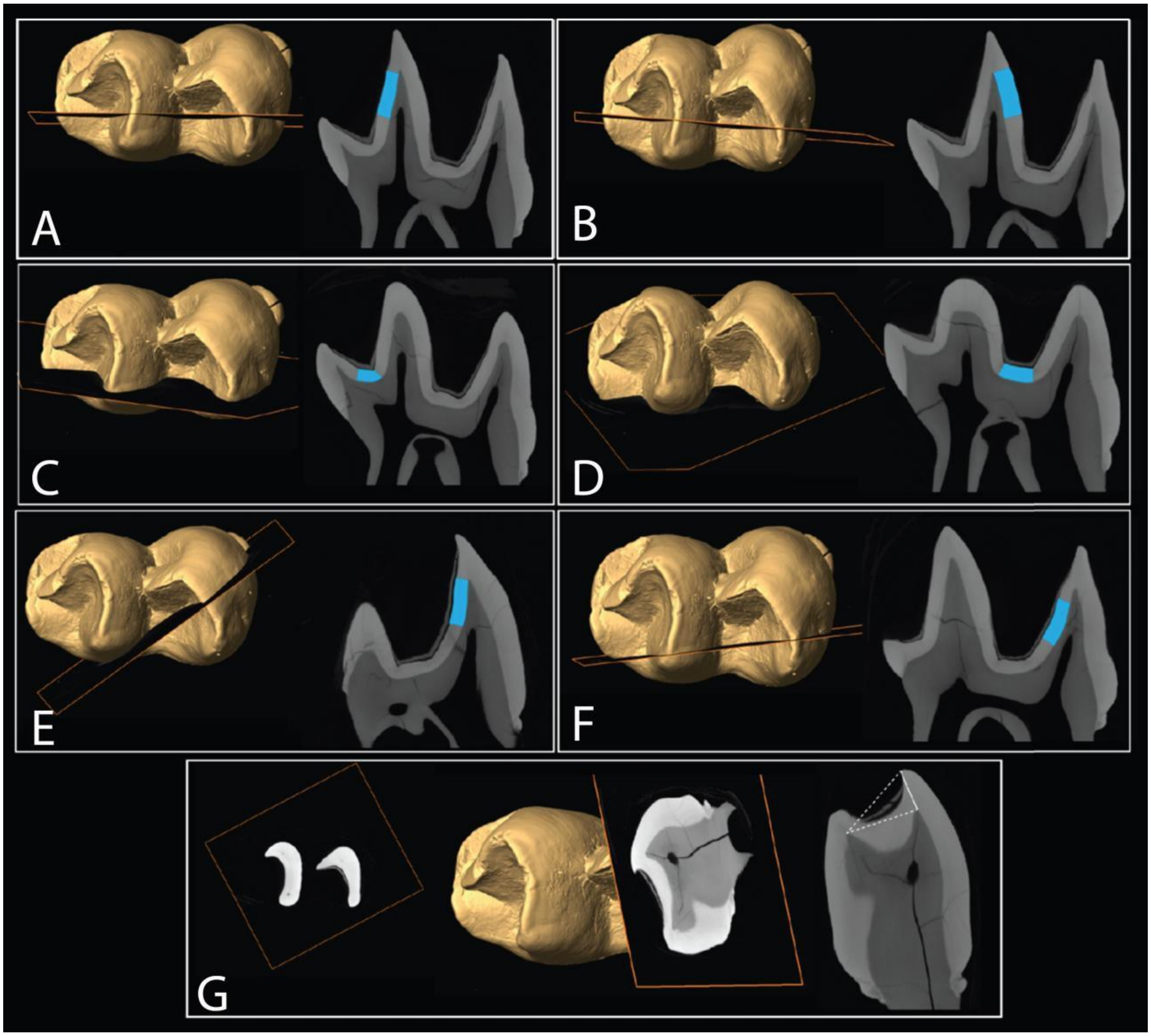


**Fig. S8. Examples of linear enamel thickness measurements on a right, lower third molar (*Osphranter rufus*, FU 2003.9.12-12).** (**A**) Enamel thickness of the anterior protolophid (AP). (**B**) Enamel thickness of the posterior protolophid (PP). (**C**) Enamel thickness of the trigonid fossa (TRGF). (**D**) The interlophid basin (INT). (**E**) Buccal face of the anterior hypolophid (AHB). (**F**) Lingual face of the anterior hypolophid (AHL). (**G**) Hypoconid apex (HCA).
